# Sex differences in contextual pattern separation, activation of new neurons, and functional connectivity within the limbic system

**DOI:** 10.1101/2022.02.28.482359

**Authors:** Shunya Yagi, Amanda Lee, Nadine Truter, Liisa AM Galea

## Abstract

Sex differences in the structure and function of hippocampus exist. Here, we examined sex differences in contextual pattern separation, functional connectivity, and activation of new neurons during fear memory. Two-month-old male and female Sprague-Dawley rats were injected with the DNA synthesis markers, iododeoxyuridine (IdU) and chlorodeoxyuridine (CldU) three weeks and four weeks before perfusion, respectively. One week after CldU injection, the rats underwent a context discrimination task in which rats were placed in context A (shock) and context A’ (no shock) every day for 12 days. On the test day, rats were placed in the shock context (context A) to measure fear memory and expression of zif268, an immediate early gene across 15 different limbic and reward regions. We found that females, but not males, showed contextual discrimination during the last days of training. On the test day, both sexes displayed similar levels of freezing, indicating equivalent fear memory for context A. Despite similar fear memory, males showed more positive correlations of zif268 activation between the limbic regions and the striatum, whereas females showed more negative correlations among these regions. Females showed greater activation of the frontal cortex, dorsal CA1, and 3-week-old adult-born dentate granular cells compared to males. These results highlight the importance of studying sex differences in fear memory and the contribution of adult neurogenesis to the neuronal network and may contribute to differences in susceptibility to fear related disorders such as posttraumatic stress disorder.

## 1. Introduction

Women are more likely to present with anxiety disorders such as post-traumatic stress disorder (PTSD) compared to men [1,2], which are associated with disrupted hippocampal integrity [3,4]. The hippocampus plays important roles for pattern separation and pattern completion [5,6]. Pattern separation is a major component of episodic memory, which refers to the process of forming distinct representations of similar inputs during memory encoding [6]. Impairments in pattern separation are involved in overgeneralization of fear memory among patients with PTSD [4,7].

Adult hippocampal neurogenesis is required for pattern separation and for stress resilience, as rodents with ablation of adult neurogenesis show impairments during pattern separation tasks and reduced stress resilience [8,9]. Studies demonstrate that hippocampusamygdala-frontal cortex connectivity plays a critical role for long-term fear memory in humans [10,11] and there are sex differences in the resting-state functional connectivity within this circuit [12]. In rodents, functional connectivity has been investigated via immediate early gene (IEG) mapping [13,14]. IEGs such as *zif268* are genes that are rapidly activated in response to neuronal stimulation and induce IEG proteins, which play an important role in neural plasticity and memory [15–17]. Brain-wide IEG imaging approaches in rodents can detect coordinated activation with high spatial resolution, which is useful to describe functional connectivity [14]. However, it remains to be determined whether new neurons in the DG are activated in a coordinated fashion with other brain regions, and whether sex modulates the functional connectivity of adult-born neurons during recall of fear memory.

Therefore, we examined sex differences in contextual pattern separation and functional connectivity among 15 different limbic and reward regions during fear memory retrieval. A fear conditioning context discrimination task was used to assess sex differences in the ability to discriminate between two contexts in male and female rats. Furthermore, a combination of thymidine analogue labelling and IEG imaging was used to examine the coordinated neuronal activation of adult-born dentate granular cells (DGC)s with other brain regions. We hypothesized that male rats would have greater ability for separating two contexts compared to females. Furthermore, we also predicted that males and females would show distinct patterns of coordinated neuronal activation of different brain regions (hippocampus, amygdala, frontal cortex and striatum) during fear memory retrieval and that new neurons would show disparate patterns of functional connectivity between the sexes.

## 2. Methods

### 2.1. Subjects

Sixteen 8-week-old Sprague Dawley rats (males: n□=□8; females: n□=□8) were purchased from Charles River Canada (St-Constant, QC, Canada). Rats were pair-housed in opaque polysulfone bins (432 mm × 264 mm × 324 mm) with paper towels, a single polycarbonate hut, virgin hardwood chip bedding, and free access to food and water. Males and females were housed in separate colony rooms that were maintained under a 12:12-h light/dark cycle (lights on at 07:00 h). All animals were handled every day for two minutes beginning one week after arrival for two weeks. All experiments were carried out in accordance with the Canadian Council for Animal Care guidelines and were approved by the animal care committee at the University of British Columbia. All efforts were made to reduce the number of animals used and their suffering during all procedures.

### 2.2. Apparatus

Behavioral testing for all experiments was conducted in four operant chambers (30.5 × 24 × 21 cm; Med-Associates, St Albans, VT) enclosed in sound-attenuating boxes which are further described in the Supplemental Methods.

### 2.3. Procedures

#### 2.3.1. Experimental timeline

Subjects received one injection of 5-chloro-2’-deoxyuridine (CldU:171 mg/kg; intraperitoneal (i.p.), MP Biomedicals, Santa Ana, CA, USA) on Experimental Day 1 and one injection of 5-iodo-2’-deoxyuridine (IdU: 56.75 mg/kg; i.p., Cayman Chemical, Ann Arbor, MI, USA) on Experimental Day 8. Thymidine analogs, IdU and CldU, incorporate into DNA during synthesis phase of cell proliferation, which can be distinguished from one another using respective antibodies [18–21]. Subjects were tested in the contextual pattern separation task (modified from [22]) for 12 days (Experimental Days 16--28, which are referred to as Trial Days 1-12), followed by a day of activation test trial that is described below (Experimental Day 29; see Figure 1A).

**Figure 1.**
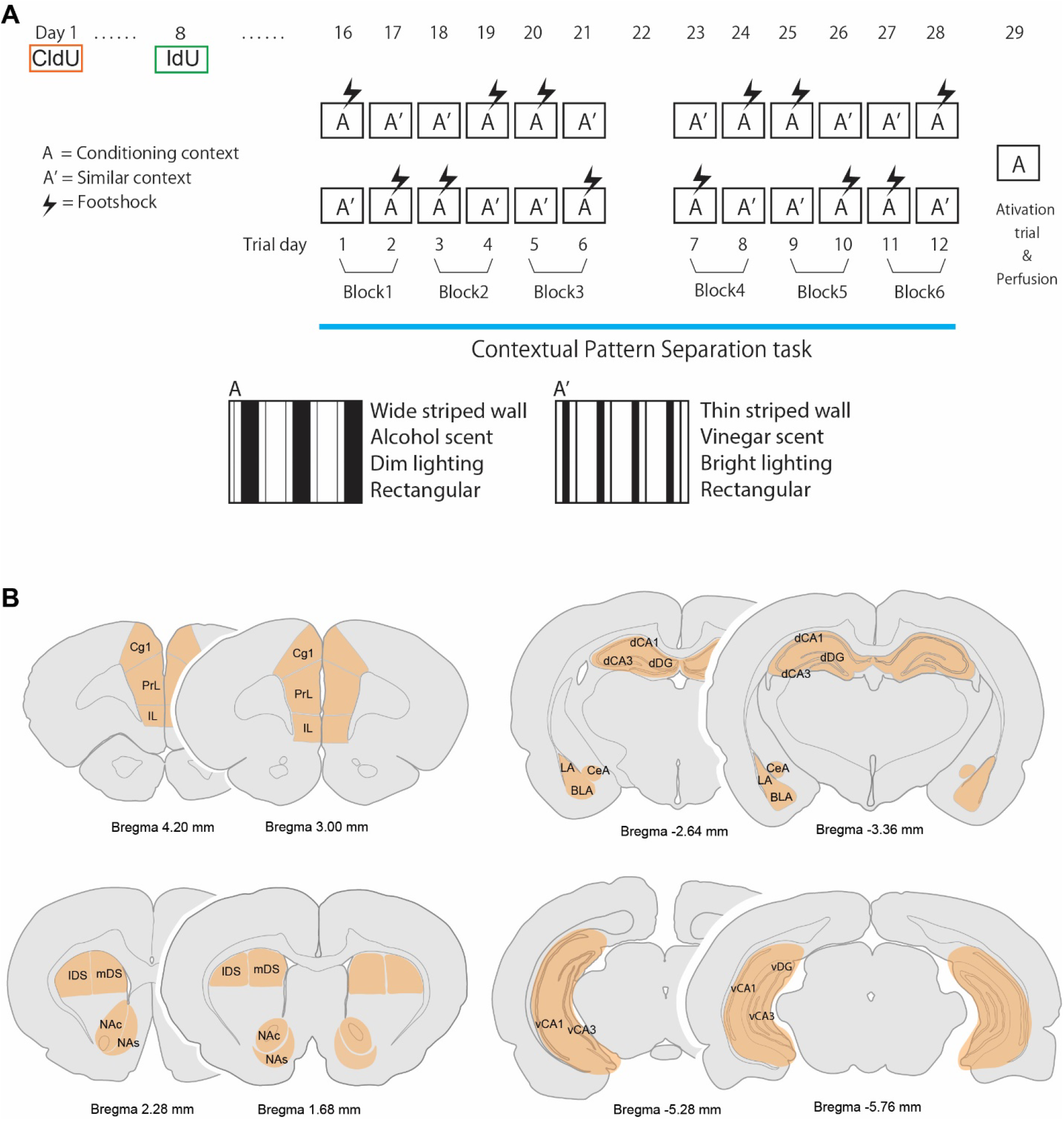
(A) Schematic illustration of experimental timeline: Subjects received one intraperitoneal injection of 5-chloro-2’-deoxyuridine on Experimental Day 1 and one intraperitoneal injection of 5-iodo-2’-deoxyuridine on Experimental Day 8. Then, subjects were tested in the contextual pattern separation task for a total of 12 days (Experimental Day 16-28), followed by an activation trial in which the rats were placed in the context previously paired with shock but received no shock (Experimental Day 29). During the contextual pattern separation task, subjects were exposed to two different contexts each day; context A a shock-paired context (context paired with foot shocks) and context A’ a neutral context (context with no foot-shock). (B) Brain regions that were examined for functional connectivity using zif268. ACC: cingulate cortex (Cg1); PrL: prelimbic cortex; IL: infralimbic cortex; lDS: lateral dorsal striatum; mDS: medial dorsal striatum; NAc: nucleus accumbens core; NAs: nucleus accumbens shell; LA: lateral amygdala; BLA: basolateral amygdala; CeA: central amygdala; dDG: dorsal dentate gyrus; vDG: ventral dentate gyrus; dCA1: dorsal cornu ammonis 1; vCA1: ventral cornu ammonis 1; dCA3: dorsal cornu ammonis 3; vCA3: ventral cornu ammonis 3.

#### 2.3.2. Behavioral testing for contextual pattern separation

Subjects were exposed daily for five minutes each to two different contexts (4-5 hours interval between contexts), a shock-paired context (Context A) and a neutral context (Context A’), for a total of 12 days. The contexts for Context A trials and Context A’ trials were counterbalanced across subjects and remained the same for each subject throughout the entire experiment. During the shock-paired trial in Context A, subjects were allowed to explore the chamber for three minutes followed by three one-second foot shocks (0.6 mA) with 30 second intervals between each shock. The subjects returned to their home cage one minute after the third shock. During the neural trial in Context A’, the subjects explored a different context from Context A for five minutes without receiving a foot shock and returned to their home cage. The order of two contexts that subjects were exposed each day for the first six days followed AA’ - A’A - A’A - AA’ - AA’ - A’A design, and the order was reversed for the remaining days [22] (see Figure 1A).

The duration of freezing during the first three minutes of each trial was examined as the conditioned fear response, and the percentage of freezing was calculated by dividing the duration of freezing by 180 seconds. A discrimination index (DI) was calculated with the following formula on the last two days of training:

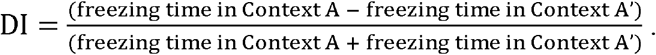

As a previous study found sex differences in darting, an active fear response, in a cued fear conditioning task [23], darting behavior was also recorded.

#### 2.3.3. Activation trial and perfusion

On the day after Training Day 12, the Activation Test Trial was conducted to examine fear memory. Subjects were exposed to the Context A for five minutes without a foot shock and returned to their home cage. Video recordings were analysed for active fear behavior (darting), passive fear behavior (freezing), or other behaviors (rearing, grooming and non-specific behaviors; see supplemental). However, no darting in our paradigm was observed. Ninety minutes after the Activation trial, subjects were administered an overdose of sodium pentobarbitol (500 mg/kg, i.p.) and perfused transcardially with 60 mL of 0.9% saline followed by 120 mL of 4% paraformaldehyde (Sigma Aldrich).

### 2.4. Tissue processing

Extracted brains were postfixed in 4% paraformaldehyde overnight, then transferred to 30% sucrose (Fisher Scientific, Ottawa, ON, Canada) solution for cryoprotection and remained in the solution until sectioning. Brains were sliced into 30-μm coronal sections using a Leica SM2000R microtome (Richmond Hill, ON, Canada). Sections were collected in series of 10 throughout the entire rostral□ caudal extent of the forebrain (Bregma 5.64 to −7.56 mm) and stored in antifreeze solution consisting of ethylene glycol, glycerol, and 0.1 M PBS at −20°C. Details of the immunohistological staining and antibodies used are found in the Supplemental Section.

### 2.5. Cell counting

All counting was conducted by an experimenter blind to the group assignment of each animal using an Olympus FV1000 confocal microscope and/or Zeiss Axio Scan.Z1 (Carl Zeiss Microscopy, Thornwood, NY, USA). Density of immunoreactive cells was calculated by dividing the total immunoreactive (ir) cells by volume (mm^3^) of the corresponding region. Volume estimates were calculated by multiplying the summed areas by thickness of sections (0.03 mm, using Cavalieri’s principle [24]). Area measurements for the region of interest were obtained using digitized images on Zen 3.0 software (blue edition; Carl Zeiss Microscopy, Thornwood, NY, USA).

Brain regions were defined according to a standard rat brain atlas (Paxinos and Watson, 2004). Location of immunoreactive cells in the hippocampus was examined in the dorsal or ventral dentate gyrus using the criterion defined by Banasr et al. (2006)[25] with sections 7.20-4.48mm from the interaural line (Bregma −1.80 to −4.52mm) defined as dorsal and sections 4.48-2.20 mm from the interaural line (Bregma −4.52 to −6.80mm) as ventral [25]. Cells were counted separately in each region because the different regions are associated with different functions (reviewed in [26]) and different maturation timelines of neurogenesis [27,28]. The dorsal hippocampus is associated with spatial reference memory, whereas the ventral hippocampus is associated with working memory, stress and anxiety [29,30]. See Supplemental Section for details on counting procedures for doublecortin, IdU and CldU.

#### 2.5.1. zif268 counting

Photomicrographs of coronal sections containing the frontal cortex, amygdala, hippocampus, striatum, nucleus accumbens were obtained from ZEISS Axio Scan.Z1 slidescanner with a 20x objective lens (four images from each region of interest: see Figure 1B). Zif268-ir cells in the infralimbic cortex (IL), prelimbic cortex (PrL), anterior cingulate cortex (ACC: Cg1), medial part of dorsal striatum (mDS), lateral part of dorsal striatum (lDS), nucleus accumbens core (NAc), nucleus accumbens shell (NAs), central nucleus of amygdala (CeA), basaolateral nucleus of the amygdala (BLA), lateral nucleus of the amygdala (LA), dorsal(d) hippocampus (dCA1, dCA3, dDG) and ventral(v) hippocampus (vCA1, vCA3, vDG) were counted automatically from the digitized images using a code developed by JEJS (see [28] for details) on MATLAB (MathWorks; Natick, Massachusetts, USA).

#### 2.5.2. Estrous-cycle determination

Daily lavage samples were taken from all females after behavioral procedures (see Supplemental Methods section). Estrous cycle determination was done as the estrous cycle stage can affect long term potentiation and IEG expression in the hippocampus [31,32]. There was one female in the proestrous stage during Activation Trial in the present study. Thus, estrous cycle phase was used as a covariate for all analyses.

### 2.6. Statistical analyses

All analyses were conducted using Statistica (Statsoft Tulsa, OK) unless otherwise stated, and significance level was set at α = 0.05. Repeated-measures or factorial analysis of variance (ANOVA), with sex (male and female) as between-subject variables were conducted on our variables of interest (freezing, zif268 expression). Post-hoc tests used the Newman-Keuls procedure. A priori comparisons were subjected to Bonferroni corrections. Effect sizes are given with Cohen’s d or partial η^2^. Pearson product-moment calculations and principal component analyses on zif268 expression across regions were also performed. Further details of the statistical procedures used are found in the Supplemental Section.

## 3. Results

### 3.1. Females, but not males, discriminated shock-paired contexts from neutral contexts

Male and female rats were exposed to 12 days of the contextual pattern separation task to examine the ability for discriminating a shock-paired context from a neutral context. Females exhibited significantly greater percentage of freezing in the shock-paired context (Context A) than in the neutral context (Context A’) on two days: Trial Day 9 (p < 0.02, Cohen’s d = 1.114), and Trial Day 12 (p < 0.0001, Cohen’s d = 1.696), whereas there were no significant differences between the contexts on any day in males (Figure 2A and 2B; interaction effect of sex by day by context [F(11, 154) = 2.25, p = 0.014, η_p_^2^ = 0.139)]). There was also a significant interaction effect of day by context [F(11, 154) = 6.26, p < 0.0001, η_p_^2^ = 0.309] and a main effect of day [F(11, 154) = 21.04, p < 0.0001, η_p_^2^ = 0.600]. Consistent with these findings, the discrimination index (DI) of the last two days (Trial Day 11 and 12 or Block 6) indicated that females showed a greater DI compared to males [F(1, 14) = 5.81, p = 0.030, Cohen’s d = 1.205; Figure 2C].

**Figure 2.**
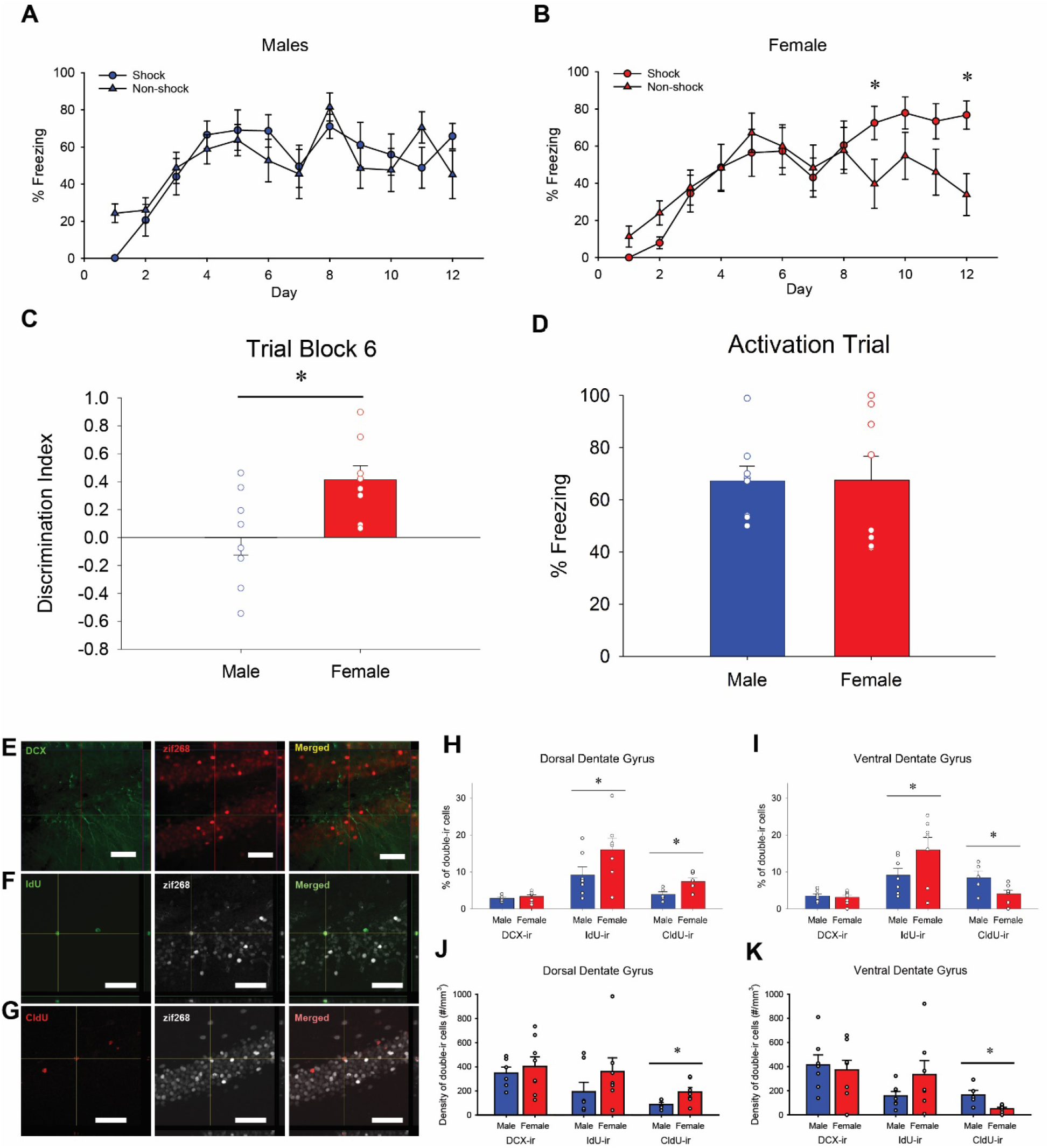
A-D: Females, but not males, discriminated shock-paired contexts from neutral contexts. Mean (±SEM) percentage of freezing in Context A (shock) and Context A’ (non-shock) in males (A) and females (B). Mean (±SEM) discrimination index of the trial block 6 (Trial Day 11+12) (C). Mean (±SEM) percentage of freezing in males and females during the activation trial on Day 29 (D). Females exhibited significantly greater percentage of freezing in the shock-paired context (Context A) than in the neutral context (Context A’) on Trial Day 9 and Trial Day 12, whereas there was no significant difference in percentage of freezing between the two contexts in any days in males. Furthermore, females showed greater discrimination based on the index on the last trial block (Trial Day 11+12) compared to males. There was no significant sex difference in the percentage of freezing during the Activation Trial. * indicates p < 0.05. **E-K: Sex differences in zif268 activation of adult-born cells in the dentate gyrus.** Photomicrographs of doublecortin-immunoreactive (ir) cells (DCX-ir: green) and zif268-ir cells (red) were taken under Zeiss Axio Scan.Z1 with 40x objective lens (E). Photomicrographs of IdU-ir cells (green) and zif268-ir cells (white) were taken under Olympus FV1000 confocal microscope with 40x objective lens (F). Photomicrographs of CldU-ir cells (red) and zif268-ir (white) cells were taken under Olympus FV1000 confocal microscope with 40x objective lens (G). Scale bars indicate 50 μm. Mean (±SEM) percentage of double-labelled cells in the dorsal (H) and ventral (I) dentate gyrus. Females, compared to males, had greater percentage of IdU/zif268-ir cells in the dorsal and ventral dentate gyrus, whereas there was a significant interaction effect of sex by region for the percentage of CldU/zif268-ir cells. Mean (±SEM) density of double-labelled cells in the dorsal (J) and ventral (K) dentate gyrus. There was a significant interaction effect of sex by region for the percentage of CldU/zif268-ir cells. * indicates p < 0.05.

On the activation trial day, all subjects were exposed to the conditioning context A (on the day after the 12 training Trials) without any shock to assess fear memory. There was no significant sex difference in the percentage of freezing during the activation trial (p = 0.932, Cohen’s d = 0.045; Figure 2D), indicating the memory strength for the shock-paired context was equivalent between the sexes, despite the better discrimination learning in females. As noted earlier, no darting behavior was observed throughout the experiment.

### 3.2. There were more 4-week-old cells in the dorsal compared to the ventral dentate gyrus

IdU and CldU were injected three weeks and four weeks, respectively, before perfusion. The density of CldU-ir cells was greater in the dDG compared to vDG [main effect of region: F (1, 11) = 6.50, p = 0.027, Cohen’s d = 1.104; see supplemental Figure 1]. There were no other significant main or interaction effects on the density of DCX-ir, CldU-ir or IdU-ir cells (p’s > 0.283).

### 3.3. Females had a greater percentage of IdU/zif268-ir cells in the dentate gyrus compared to males

The percentage of DCX-ir, IdU-ir, or CldU-ir cells that were double-labelled with zif268 was measured to examine neural activation of adult-born cells in the DG. Females, compared to males, had a greater percentage of IdU/zif268-ir cells in the DG [main effect of sex: F (1, 12) = 4.59, p = 0.05, Cohen’s d = 1.539; Figure 2H and 2I] but no other main or interaction effects for IdU/zif268-ir cells. Females, compared to males, had greater percentage of CldU/zif268-ir cells in the dDG (p = 0.047, Cohen’s d = 1.538), whereas, males, compared to females, had greater percentage of CldU/zif268-ir cells in the vDG (p = 0.015, Cohen’s d = 1.317) [interaction effect of region by sex: F(1, 10) = 19.53, p = 0.001, η_p_^2^ = 0.661; Figure 2H and 2I]. There were no significant main effects (all p’s > 0.46) for CldU/zif268-ir cells. There were no significant effects of DCX/zif268-ir cells (p’s > 0.66).

Along with the percentage of double-labelled cells we also examined the density of DCX-ir, IdU-ir, and CldU-ir cells that were double-labelled with zif268. Females, compared to males, had greater density of CldU/zif268-ir cells in the dDG (p = 0.033), whereas, males, compared to females, had greater density of CldU/zif268-ir cells in the vDG (p = 0.023) [interaction effect of region by sex: F(1, 10) = 19.34, p = 0.001, η_p_^2^ = 0.659; Figure 2J and 2K]. There were no main or interaction effects for the density of IdU/zif268-ir cells or DCX/zif268-ir cells (all p’s > 0.106).

### 3.4. Females had greater zif268 immunoreactivity than males in the frontal cortex and dorsal CA1 region of the hippocampus in response to a shocked-paired context

The density of zif268-ir cells was measured to examine neural activation in subregions of the frontal cortex, dorsal striatum, nucleus accumbens, hippocampus and amygdala in response to exposure to the shocked-paired context. Females, compared to males, showed greater density of zif268-ir cells across the different regions of the frontal cortex [main effect of sex: F(1, 12) = 10.14, p = 0.008, η_p_^2^ = 0.458], as well there was greater density of zif268-ir cells in the ACC and PrL compared to IL [main effect of subregion: F(2, 24) = 34.40, p < 0.001, η_p_^2^ = 0.741; post-hoc: all p’s < 0.001; Figure 3C].

**Figure 3.**
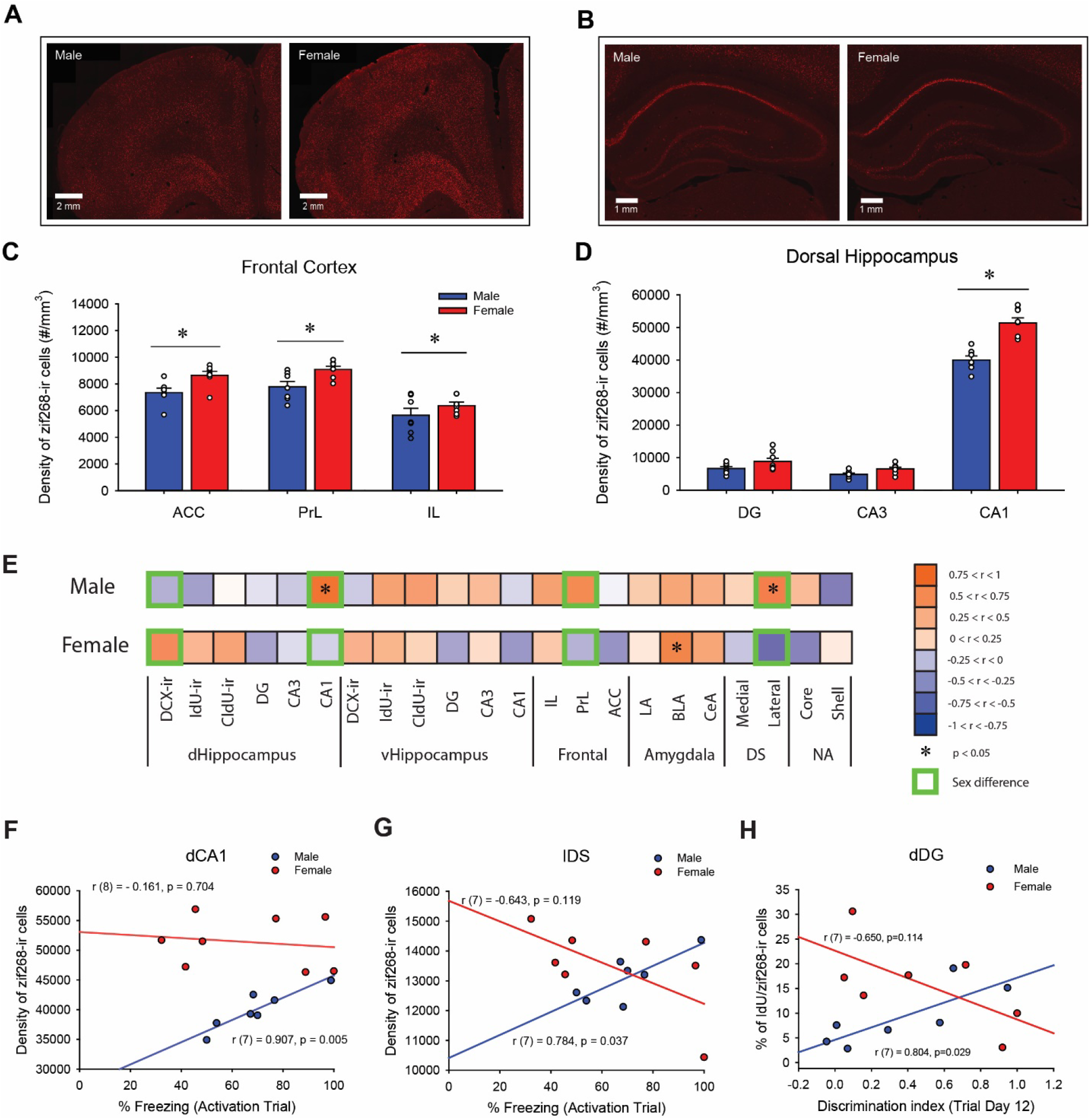
A-D: Sex differences in zif268 activation in the brain in response to the shock-paired context. Photomicrographs of zif268 immunoreactivity in the frontal cortex (A) and in the dorsal hippocampus (B) in male (left) and female rats (right). Mean (±SEM) density of zif268-ir cells in the frontal cortex (C) and in the dorsal hippocampus (D). Females, compared to males, had greater density of zif268-ir cells in the anterior cingulate cortex (ACC), in the prelimbic cortex (PrL) and in the dorsal CA1. * indicates p < 0.05. **E-H: Sex differences in correlations between the amount of freezing during the activation trial and zif268 activation across 22 brain regions.** Heat maps generated based on correlations coefficients (E). Males and females had significant correlations of zif268-ir cell density in different brain regions with the percentage of freezing (* indicates correlations with p < 0.05). There were significant sex differences in the correlations between the percentage of freezing and the density of zif268-ir cells in four different brain regions (Green boxes indicates significant sex differences with p < 0.05). Scatter plots for correlations between the percentage of freezing and the density of zif268-ir cells in the dorsal CA1 (dCA1) (F) or in the lateral dorsal striatum (lDS) (G), and correlations between the discrimination index on the last trial day and the percentage of IdU/zif268-ir cells in the dorsal dentate gyrus (H) in males (blue) and females (red).

Furthermore, females showed greater density of zif268-ir cells in the dorsal CA1 compared to males [p < 0.001, Cohen’s d = 2.957; interaction effect of sex by subregion by dorsoventral axis: F(2, 26) = 17.56, p < 0.001; Figure 3D]. In both males and females, the density of zif268-ir cells in the dCA1 is greater than dDG and dCA3, and the density of zif268-ir cells in the vCA1 is greater than vCA3 (all p’s < 0.01). There were also significant interaction effects of sex by subregion [F(2, 26) = 10.36, p < 0.001, η_p_ = 0.444], sex by dorsoventral axis [F(1, 13) = 10.82, p = 0.006, η_p_^2^ = 0.454] and subregion by dorsoventral axis [F(2, 26) = 1028.60, p < 0.001, η_p_^2^ = 0.988], and main effects of sex [F(1, 13) = 21.53, p < 0.001, η_p_^2^ = 0.624], subregion [F(2, 26) = 819.06, p < 0.001, η_p_^2^ = 0.984] and dorsoventral axis [F(1, 13) = 690.17, p < 0.001, η_p_^2^ = 0.982].

There were no significant main or interaction effects involving sex in activation in the amygdala, striatum or the nucleus accumbens, but there were significant regional differences within these areas. The lDS had greater density of zif268-ir cells compared to the mDS [main effect of subregion: F(1, 12) = 6.88, p = 0.022, η_p_^2^ = 0.364; supplemental figure 2A]. In the amygdala, the LA had greater density of zif268-ir cells compared to the CeA (p = 0.007) and the BLA (p < 0.001), and the CeA had greater density of zif268-ir cells compared to the BLA (p < 0.001) [main effect of subregion: F(2, 24) = 28.45, p < 0.001, η_p_^2^ = 0.703; Supplemental Figure 2B]. In the nucleus accumbens, the density of zif268-ir cells was significantly greater in the shell compared to the core [main effect of subregion: F(1, 12) = 22.22, p < 0.001, η_p_^2^ = 0.649; Supplemental Figure 2C]. There were no significant main or interaction effects of sex for the density of zif268-ir cells in any of these regions (P > 0.111).

### 3.5. The density of zif268-ir cells in the dorsal CA1 was positively correlated with amount of freezing during memory recall in males, but not in females

Pearson product-moment correlations were calculated between the percentage of freezing during the activation trial and the density of zif268-ir cells in the 16 brain regions and the six different populations of adult-born cells (dorsal or ventral DCX/IdU/CldU co-expressing zif268; see Figure 3E). There was a significant positive correlation between the density of zif268-ir cells and the percentage of freezing in males in the dCA1 [r (7) = 0.907, p = 0.005; Figure 3F]. There were no other significant correlations between the density of zif268-ir cells and the percentage of freezing during the activation trial after Bonferroni corrections (p’s > 0.037). Sex differences in the correlations were noted, with males having positive correlations and females having negative correlations, between the percentage of freezing and the density of zif268 cells in the dDG (p = 0.041), dCA1 (p = 0.006; Figure 3F), PrL (p =0.040) and lDS (p = 0.005; Figure 3G). See Supplemental Results for details.

### 3.6. Neural activation of adult-born cells in the dentate gyrus was associated with the ability for pattern separation in males but not females

Pearson product-moment correlations were calculated between the percentage of DCX, IdU or CldU-ir cells that were double-labelled with zif268 and the discrimination index (DI) on the last trial day. The percentage of IdU-ir cells that were double-labelled with zif268 in the dDG was significantly correlated with DI in males [r(7) = 0.804, p = 0.029], but not in females [r(7) = −0.650, p = 0.114; Figure 3H], which was significantly different between the sexes (p = 0.004). There were no other significant correlations between the percentage of DCX, IdU or CldU-ir cells that were double-labelled with zif268 and the DI (all p’s > 0.127).

### 3.7. Males and females showed distinct patterns of significant inter-regional correlations of the density of zif268-ir cells

Pearson product-moment correlations were calculated with the density of zif268-ir cells between 16 brain regions and six different populations of adult-born cells (double-labelled with zif268 and DCX, IdU, or CldU in the dorsal or ventral DG) to examine functional connectivity between these regions to activated new neurons of different ages (see Figure 4). As can be seen in Figure 4B and 4C there were mainly positive correlations between activation of new neurons of different ages within the hippocampus in males (19), with much fewer seen in females (4). In addition, there were more correlations of activated new neurons with regions outside the hippocampus in females (7, with only 2 significant to the amygdala) than in males (4 with none significant) (see Supplemental Table 1 for the detailed statistical data). The Fischer z-test statistic revealed significant sex differences in the 27 inter-regional correlations (see Supplemental Results for the detailed statistical data).

**Figure 4.**
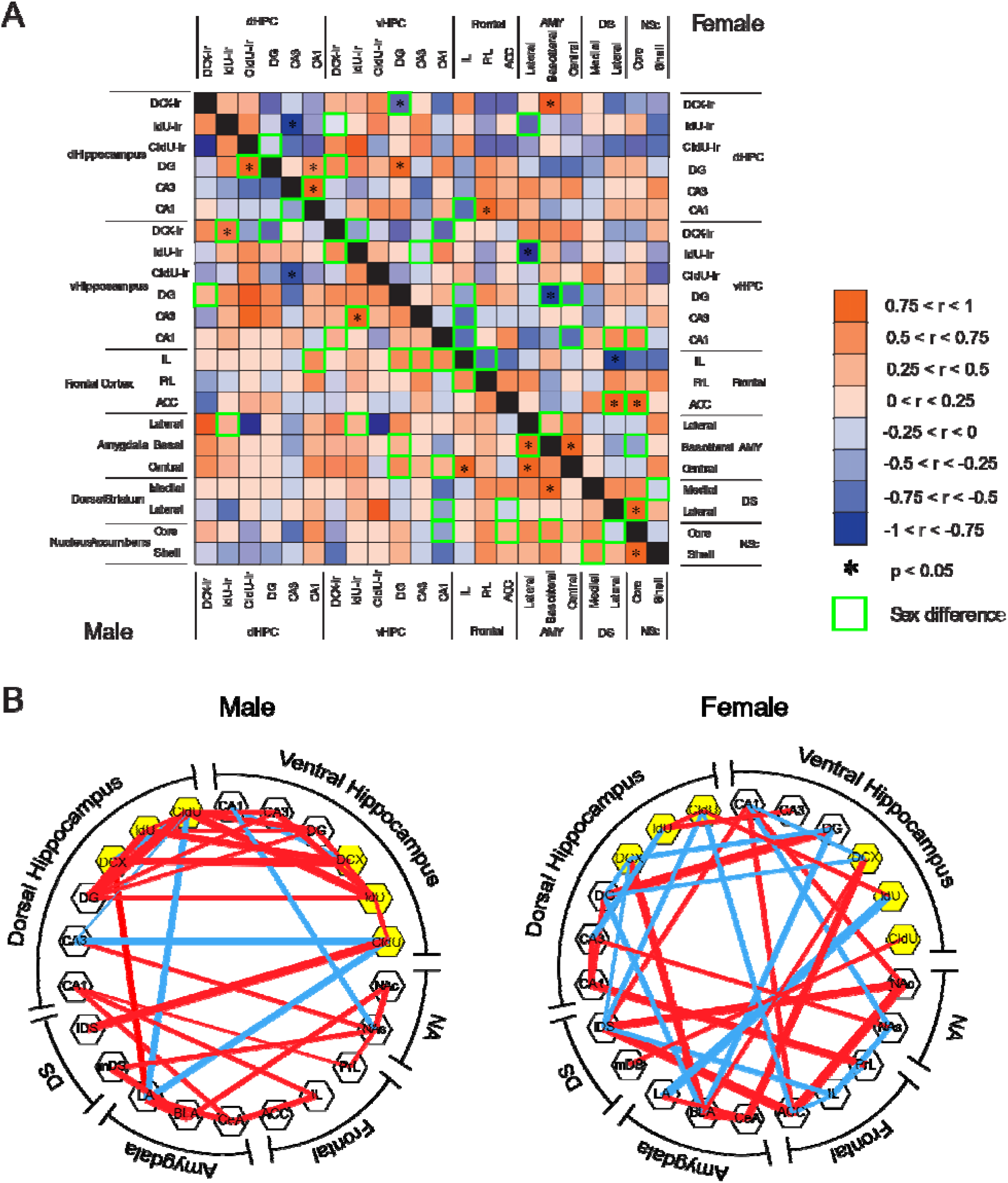
Sex differences in inter-regional correlations of zif268-ir cell density. **A:** A heatmap showing correlation coefficients (r) of the density of zif268-ir cells between each brain region in males and females. Males and females showed distinct patterns of significant inter-regional correlations of zif268-ir cell density. * indicates significant correlations (p < 0.05) and green boxes indicate sex differences between the correlations (p < 0.05). **B:** Brain network maps were generated with correlations with coefficients larger than 0.67 or smaller than −0.67 in males (left) and females (right) with p<0.1. Red lines indicate positive correlations with wider lines indicating larger coefficients and blue lines indicate negative correlations with wider lines indicating smaller coefficients.

### 3.8. Principal component analyses on the density of zif268-ir cells

Principal component analyses (PCA) were conducted with the density of zif268-ir cells in the 16 brain regions. PCA demonstrated that the first three principal components factors accounted for 70.45% of the variance with PC1 explaining 37.50% of the variance, PC2 explaining 18.57% and PC3 explaining 14.38% of the variance (see Figure 5B). PC1 included significant positive loading on the density of zif268-ir cells in all of the hippocampus, most of the frontal cortex and the amygdala (except the IL and BLA) and included the mDS (see Figure 5A). A repeated-measures ANOVA on the principal component scores revealed that females showed significantly greater positive scores compared to males in PC1 [interaction effect of sex by factor: F (2, 24) = 9.11, p = 0.001; post-hoc: p = 0.031; see Figure 5C], indicating that females had greater activation of zif268 among these regions compared to males.

**Figure 5.**
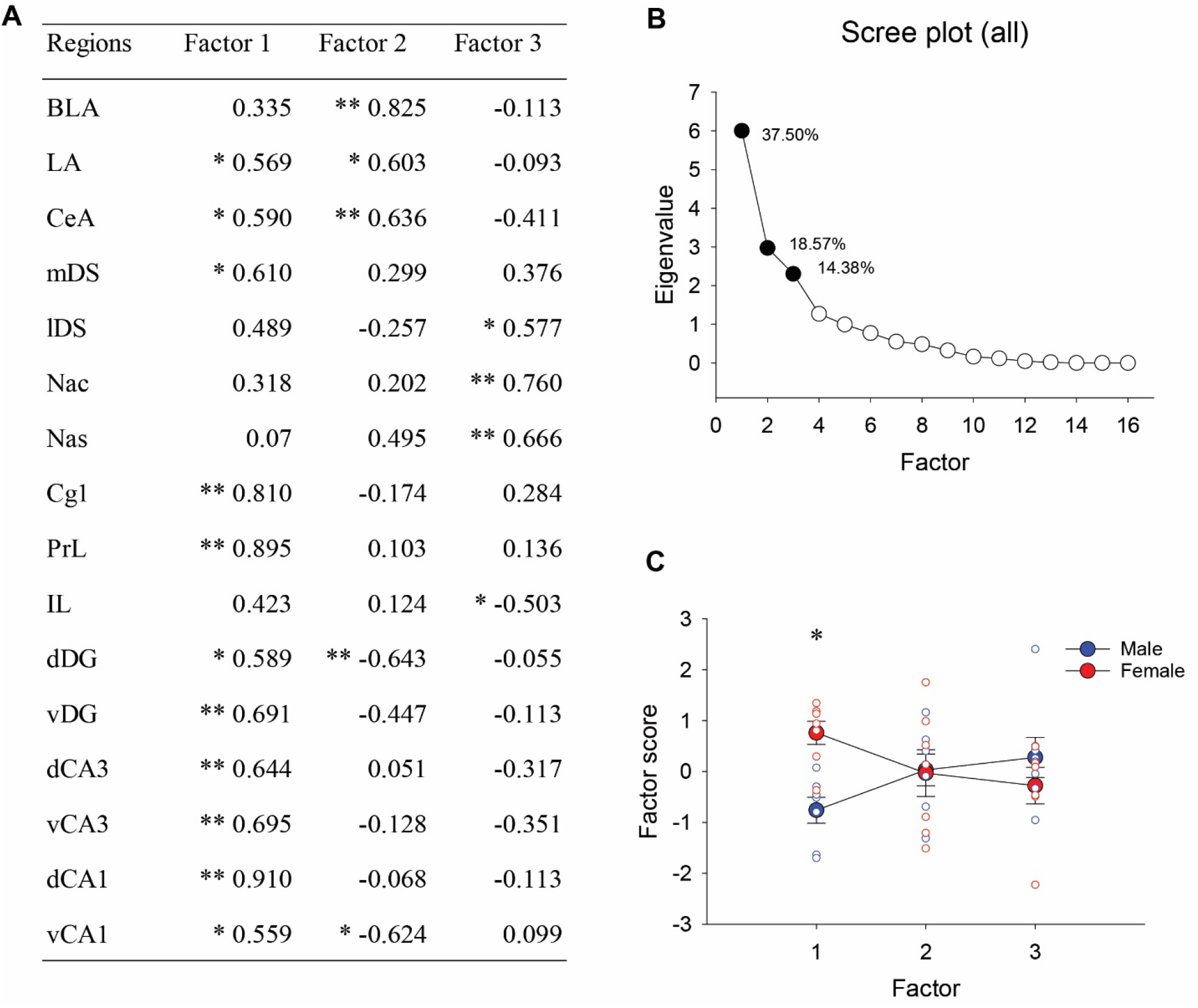
A-C: Sex difference in the principal component analysis (PCA). Factor coordinates of the variables in the first three factors identified using principal component analysis (PCA) for zif268 activation (A). Scree plot for eigenvalues with the total percentage of variance for the first three factors (B). Eigenvalues for the first three factors were significant based on Horn’s parallel analysis. The first three factors explained 70.45% of the variances. A graph showing sex difference in the factor scores of individual samples (C). ANOVA and post-hoc revealed males and females showed significant sex difference in the first factor. * indicates p < 0.05 and ** indicates p < 0.01.

## 4. Discussion

We found that female rats were better at contextual pattern separation and had greater neural activation in the frontal cortex, dorsal CA1 region and in adult-born dentate granule cells (DGCs), in response to fear memory compared to males. Furthermore, we found distinct sex differences in functional connectivity in both valence and in activation patterns between limbic regions, as males had more positive correlations among these regions than females. Intriguingly, we saw that activation of new neurons of different ages were intercorrelated and correlated with different regions in the hippocampus in males but that females, but not males, showed significant correlations between activated new neurons and the amygdala during fear memory retrieval. These results demonstrate that males and females employ different brain networks during fear memory retrieval. These findings highlight the importance of studying sex differences in fear memory and the contribution of adult neurogenesis to the neuronal network. They also have implications for targeting treatment of fear-related disorders between males and females.

### 4.1. Females are better at fear-associated contextual pattern separation compared to males

Females showed greater discrimination of the two contexts compared to males on the last trial days using the discrimination index. This result is consistent with previous studies that reported a female advantage in fear-conditioning context discrimination tasks in rodents [33,34] and in performance in emotional episodic memory tasks in humans [35–37]. However, others have found the opposite, with a male advantage in fear-conditioning context discrimination that used different spatial configurations for the contexts [38]. Indeed, in our own previous work, we found a male advantage in the ability for spatial pattern separation in rats [39], that was observed only among rats that rely more on allocentric (geometric) spatial cues [39]. The inconsistency between findings across studies may be due to sex differences in learning strategies and the types of cues between the paradigms. Males rely preferentially on spatial strategies whereas females rely more idiothetic strategies in human and rodents [40–43], which may explain why a male advantage was found when using different geometry (or shapes) of the conditioning chambers [38]. In contrast, the two contexts in the present study shared the same geometric cues, making it more difficult for rats relying on geometric cues to discriminate between the two contexts. Together these results suggest that males and females process contextual information differently and/or males and females rely on different learning strategies during a fear-conditioning context discrimination task so that the availability of favored memory cues influences their performance during a given task [44,45]. Further research is needed to determine whether there are sex differences in strategy use during a contextual learning and to determine what types of memory cues contribute to the sex difference in the ability for contextual pattern separation.

### 4.2. Females show greater neuronal activation of young granule cells in the dorsal DG compared to males

We found that 3-week-old adult-born DGCs showed greater neuronal activation in females compared to males in response to fear memory retrieval. Our previous work demonstrates that adult-born DGCs in male rats mature faster than in females [28]. Therefore, it is possible that female 3-week-old adult-born DGCs are more immature and highly excitable in response to fear memory retrieval compared to males, however then we might have expected to see a sex difference favoring males in activation of the mostly younger DCX-ir cells which was not the case. Another possible explanation for the sex difference in neural activation of 3-week-old adult-born DGCs is that pattern separation circuits are differently recruited during reexposure to familiar environment in female compared to male rats. Indeed, we did see different patterns of activation of activated new neurons in females compared to males, with females showing coordinated activation of new neurons with the amygdala whereas in males there were more intercorrelations within the hippocampus. Previous studies demonstrated that adult-born DGCs play different roles depending on the age of DGCs, as younger DGCs play a role for pattern separation while older DGCs play a role for pattern completion [8,46]. Therefore, greater neural activation of younger DGCs in females during memory retrieval may indeed be the reason for superior pattern separation performance by female rats compared to male rats in this task.

In addition to 3-week-old DGCs, we found sex differences in neural activation of 4-week-old DGCs depending on its location along the longitudinal axis. Females exhibited greater neural activation of 4-week-old adult-born DGCs in the dorsal DG whereas males exhibited greater neural activation of 4-week-old adult-born DGCs in the ventral DG compared to the opposite sex. This result suggests that 4-week-old adult-born DGCs play different functional roles in the dorsal and ventral DG between males and females, or males and females differently recruit 4-week-old adult-born DGCs in the dorsal and ventral DG during contextual fear conditioning paradigms. The dorsal hippocampus plays an important role for spatial learning and memory, and the ventral hippocampus is important for regulation of stress [29,30,47,48]. However, sex differences in the contribution of adult-born DGCs depending on its location along the longitudinal axis and depending on maturity of DGCs to the hippocampal cognition have yet to be determined.

### 4.3. Females show greater neuronal activation in the frontal cortex and dorsal CA1 in response to fear memory retrieval

We found that females, compared to males, showed greater neural activation in the frontal cortex despite there being no significant sex difference in fear memory during the activation trial. Previous studies have demonstrated that females have greater reliance on the frontal cortex (PrL, IL) to auditory fear memory acquisition, extinction and recall [49–52]. Collectively, these studies suggest that females rely on the frontal cortex to maintain fear memory.

Females also showed greater neural activation in the dorsal CA1 in response to fear memory than in males, consistent with other studies in contextual fear retrieval [53]. The CA1 in the hippocampus plays important roles for pattern completion during memory retrieval [54]. Further research is warranted to elucidate sex differences in the functional roles of dorsal CA1 during various memory tasks.

### 4.4. Males and females show distinct patterns of functional connectivity between frontal cortex, hippocampus and amygdala

The present study indicates significant sex differences in functional connectivity between the amygdala (LA, CeA), hippocampus (all subregions), dorsal striatum (mDS) and frontal cortex (PrL, ACC) where females show stronger positive connectivity among the regions in response to fear memory retrieval. This finding is consistent with resting-state functional connectivity and BOLD-signal changes in response to fear conditioning in humans [12,55,56] and with functional connectivity in rats [57]. Human females have greater resting-state functional connectivity between the amygdala, frontal regions and the hippocampus than human males [12,55] and show greater BOLD-signal changes in the amygdala, and anterior cingulate cortex compared to males to fear-conditioned stimuli in humans [56]. Furthermore, we found sex differences in patterns of associations between neural activation in adult-born DGCs and neural activation in other brain regions, with females showing correlations to the amygdala that were not seen in males. To our knowledge, this is the first study demonstrating sex differences in functional connectivity of adult-born DGCs and other brain regions. Overall, our study indicated significant involvement of the hippocampus in the functional connectivity in the present study, more so in females compared to males.

## Conclusion

Our data demonstrate that females, compared to males, show greater context discrimination, greater activation of 3-week-old adult-born DGCs in response to memory retrieval, and strong functional connectivity in the frontal cortex, the hippocampus, dorsal striatum and the amygdala during fear memory retrieval. Our findings indicate that sex differences exist in the underlying neural mechanisms and network activation even when no significant sex difference is observed in fear memory retrieval. Our work highlights the importance of elucidating sex-specific neural connections that may contribute to differences in susceptibility to fear related disorders such as PTSD. It also underscores that any treatments for fear-related disorders will need to consider sex as very different neural mechanisms may be underlying fear memory.

## Funding and disclosure

This work was supported by a Natural Sciences and Engineering Research Council (NSERC) Discovery Grant to LAMG, Killam Doctoral Award and Djavad Mowafaghian Center for Brain Health Endowment Award to SY. The authors have nothing to disclose.

## Acknowledgment

We would like to thank Jared Splinter, Yanhua Wen and Stephanie Lieblich for the exceptional technical assistance with this work.

## Author contributions

**SY:** Conceptualization, Methodology, Data curation, Writing-Original draft preparation, Visualization. **AL**: Visualization, Investigation. **NT**: Software, Validation. **LG**: Conceptualization, Methodology, Analysis, Writing-Reviewing and Editing, Supervision.

## Supplemental Section

### Methods

#### Apparatus

The boxes were equipped with a fan to provide ventilation and to mask extraneous noise. All behaviors were monitored and recorded by a single video camera mounted on the ceiling of each box. The chambers were equipped with a single 100-mA houselight located in the top center of a wall and the chamber floor consisted of 23 metal grid bars (0.4 cm in diameter) that ran parallel to the shorter wall of the chamber, which connected to a shock generator. Two chambers had wide vertical black (18 mm width) and white (12 mm width) stripe patterns on the walls and wiped with vinegar before and after each animal. The other two chambers had narrow vertical black (12 mm width) and white (12 mm width) stripe patterns on the walls and wiped with 70% isopropanol before and after each animal (see Figure 1). All the chambers were connected to a computer through a digital interface that recorded all experimental settings.

#### Immunohistochemistry

Brain tissue was double-stained for the immature neuronal protein, doublecortin (DCX), and the immediate early gene, zif268 (Figure 2A). A majority (70% or more) of adult-born granule cells express DCX within 24 hours after mitosis for up to two weeks, with maximal expression at 4 days after mitosis, and that DCX expression is rapidly reduced three weeks after mitosis (less than 20%) in both male and female rats [28,58,59]. Therefore, we used DCX to examine a cell population of new neurons that were larger 2 weeks old or younger. In addition, tissue was triple-stained for IdU, CldU, and zif268 to examine neural activation of 3-week-old (IdU) cells and 4-week-old (CldU) cells in the dentate gyrus (Figure 2B and 2C).

#### Doublecortin/zif268 double labeling

Free-floating sections were prewashed three times for 10 minutes with 0.1 M TBS. Sections were then incubated in a primary antibody solution containing 1:500 rabbit anti-zif268 (Santa Cruz Biotechnology, Dallas, TX, USA), 1:500 goat anti-doublecortin (Santa Cruz Biotechnology, Dallas, TX, USA) 0.3% Triton-X and 3% NDS in 0.1 M TBS for 24 hours at 4 °C. Sections were washed three times for 10 minutes in TBS and a further incubation of sections commenced in a secondary antibody solution containing 1:500 donkey anti-rabbit ALEXA 594 (Invitrogen, Burlington, ON, Canada), 1:500 donkey anti-goat ALEXA 488 (Invitrogen, Burlington, ON, Canada), 3% NDS and 0.3% Triton-X in 0.1 M TBS for 24 hours at 4 °C. Following three final rinses with TBS, the sections were mounted onto microscope slides and cover-slipped with PVA DABCO.

#### IdU/CldU/zif268 triple labeling

Two different thymidine analogues (CldU and IdU) were visualised with CldU-specific (rat monoclonal, clone BU1/75) and IdU-specific (mouse monoclonal, clone B44) antibodies (Podgorny et al., 2018), coupled with labelling using the immediate early gene, zif268 antibody (rabbit polyclonal). Briefly our protocol was as follows: free-floating sections were prewashed three times for 10 minutes with 0.1 M tris buffer saline (TBS; Sigma-Aldrich, Oakville, ON, Canada). Sections were then incubated in a primary antibody solution containing 1:500 rabbit anti-zif268 (Santa Cruz Biotechnology, Dallas, TX, USA), 0.3% Triton-X (Sigma-Aldrich) and 3% normal donkey serum (NDS; MilliporeSigma, Burlington, MA, USA) in 0.1 M TBS for 24 hours at 4 °C. Next, sections were incubated in a secondary antibody solution containing 1:250 donkey anti-rabbit ALEXA 647 (Invitrogen, Burlington, ON, Canada), 0.3% Triton-X, and 3% NDS in 0.1 M TBS, for 18 hours at 4 °C. After being rinsed three times for 10 minutes with TBS, sections were washed with 4% paraformaldehyde for 10 minutes, and rinsed twice in 0.9% NaCl for 10 minutes, followed by incubation in 2N HCl (Fisher Scientific, Waltham, Massachusetts, USA) for 30 minutes at 37 °C. Sections were then rinsed three times in TBS for 10 minutes each and incubated in a CldU primary antibody solution consisting of 1:1000 rat anti-BrdU (BU1/75; Abcam; Toronto, ON, Canada), 3% NDS, and 0.3% Triton-X in 0.1 M TBS for 24 hours at 4 °C. Sections were then incubated in an IdU primary antibody solution consisting of 1:500 mouse anti-BrdU (B44; BD Biosciences, San Jose, CA, USA), 0.3% NDS, and 0.3% Triton-X in 0.1 M TBS for 24 hours at 4 °C. Sections were then washed twice for 10 minutes each in a high stringency wash solution consisting of 32mM tris buffer, 50mM NaCl and 0.5% tween (pH 8.0) at 37 °C. Following three washes in TBS, sections were incubated in a secondary antibody solution containing 1:500 donkey anti-rat ALEXA 594 (Invitrogen, Burlington, ON, Canada), 1:500 donkey anti-mouse ALEXA 488 (Invitrogen, Burlington, ON, Canada), 3% NDS and 0.3% Triton-X in 0.1 M TBS for 24 hours at 4 °C. Following three final rinses with TBS, the sections were mounted onto microscope slides and cover-slipped with PVA DABCO.

#### IdU and CldU counting

Thymidine analogue immunoreactive (IdU-ir and CldU-ir) cells were counted under a 40x objective lens using Olympus FV1000 confocal microscopy. Every 20th section of the granule cell layer (GCL) that includes the subgranular zone (SGZ) was counted. The SGZ was defined as a narrow layer of cells within 30μm (equivalent to the width of three granule cell bodies) away from the innermost edge of GCL (Redila and Christie, 2006).

The percentages of IdU/zif268 and CldU/zif268-ir cells were obtained by randomly selecting 200 IdU-ir or 200 CldU-ir cells (100 cells from dorsal and 100 cells from ventral DG) and calculating the percentage of cells that were double-labelled with zif268 under a 40x objective lens using Olympus FV1000 confocal microscopy. Density of DCX/zif268-ir, IdU/zif268-ir or CldU/zif268-ir cells were calculated by multiplying the density of IdU-ir or CldU-ir cells by the percentage of double-labelled cells.

#### Doublecortin counting

Doublecortin immunoreactive (DCX-ir) cells were counted on digitized images on Zen 3.0 software (blue edition). Photomicrographs were taken from four dorsal and four ventral hippocampi using a ZEISS Axio Scan.Z1 slidescanner with a 40x objective lens. The percentages of DCX/zif268-ir cells were obtained by randomly selecting 200 DCX-ir cells (100 cells from dorsal and 100 cells from ventral DG) and calculating the percentage of cells that were double-labelled with zif268 on Zen 3.0 software. Density of DCX/zif268-ir cells were calculated by multiplying the density of DCX-ir cells by the percentage of DCX/zif268-ir cells. Estrous Cycle Determination. Vaginal cells suspended in water were obtained using a glass pipette, transferred onto microscope slides, stained with Cresyl Violet (Sigma), and analyzed using a 20× objective. Proestrous stage was determined when 70% of the cells were nucleated epithelial cells [60].

#### Data Analysis

The percentage of freezing during the Context A trials and Context A’ trials in the contextual pattern separation task was analyzed using repeated-measures analysis of variance (ANOVA), with sex (male and female) as between-subject variables and context (Context A and Context A’) and trial day (1^st^ – 12^th^ day) as within-subject factors. The discrimination index of the last trial block and percentage of freezing during the activation trial were analyzed using one-way ANOVA with sex as between-subject variable. The density of adult-born cells (DCX-ir, IdU-ir or CldU-ir cells) and those double-labelled with zif268 in the dentate gyrus were each analyzed using repeated-measures ANOVA with sex as between-subject variable and region (dorsal and ventral) as within-subject variable. The density of zif268-ir cells in each region (frontal cortex, dorsal striatum, nucleus accumbens, amygdala) was analyzed separately using repeated-measures ANOVA with sex as between-subject variables and subregions (frontal cortex: IL, PrL, ACC; dorsal striatum: lateral, medial; nucleus accumbens: core, shell; amygdala: central, lateral, basal; hippocampus: dorsal and ventral CA1, CA3 and DG) as within-subject variables.

Pearson product-moment correlations between the percentage of freezing and the density of zif268-ir cells were calculated in the regions of interest. For functional connectivity, Pearson product-moment correlations were calculated with the density of zif268-ir cells between each brain region. To examine the functional connectivity of adult-born cells in the dentate gyrus with the other brain regions, correlations were also calculated between the density of IdU/zif268-ir, CldU/zif268-ir or DCX/zif268-ir cells and the density of zif268-ir cells in each region. Interregional correlations were compared between the two sexes (male and female) using the single-sided observed Fischer z-test statistic.

Principal component analyses were conducted to assess brain networks that explain variances of zif268-ir cell density in the regions of interest. PCA data analyses were conducted using Statistica and R (3.4.3) statistical analysis software with the “FactoMineR” package. Horn’s parallel analysis was used to determine which component factors were retained for further analyses [61]. Horn’s parallel analysis was conducted using R (3.4.3) statistical analysis software with the “psych” package. Following the PCA, repeated-measures ANOVA was conducted to analyze principal component scores for individual samples with sex (male, female) and factors (1^st^, 2^nd^, 3^rd^) as the within-subject variable and sex (male, female) as the between-subject variable.

**Supplemental Table 1:** Mean (±SEM) duration of rearing, grooming and non-specific behaviors in males and females during the activation trial. There were no significant sex differences in any of the behaviors (p > 0.889).

|  | Behaviors (sec) |  |  |
| --- | --- | --- | --- |
|  | Rearing | Grooming | Other |
| Male | 4.38±2.03 | 0 | 54.5±8.81 |
| Female | 3.38±1.51 | 1.50±1.13 | 55.75±16.08 |

**Supplemental Figure 1.**
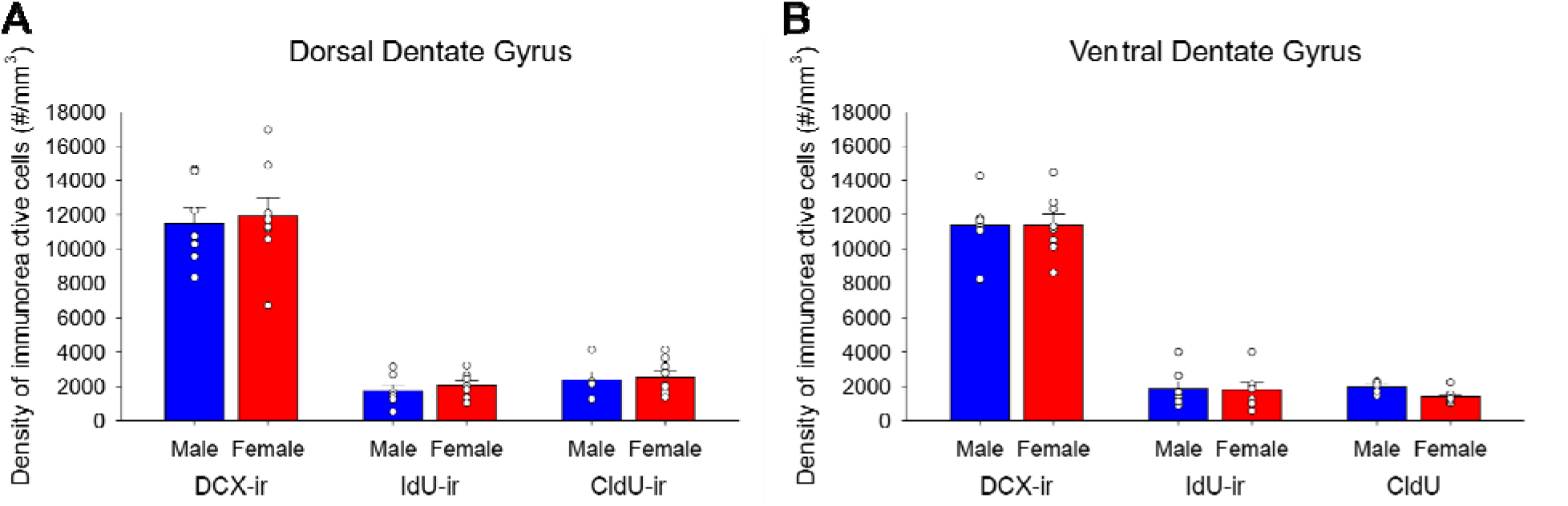
A-B: Mean (±SEM) density of adult-born cells in the dorsal (A) and ventral (B) dentate gyrus. There were no significant sex differences in the density of DCX-ir cells, IdU-ir cells or CldU-ir cells in the dorsal or ventral dentate gyrus.

**Supplemental Figure 2.**
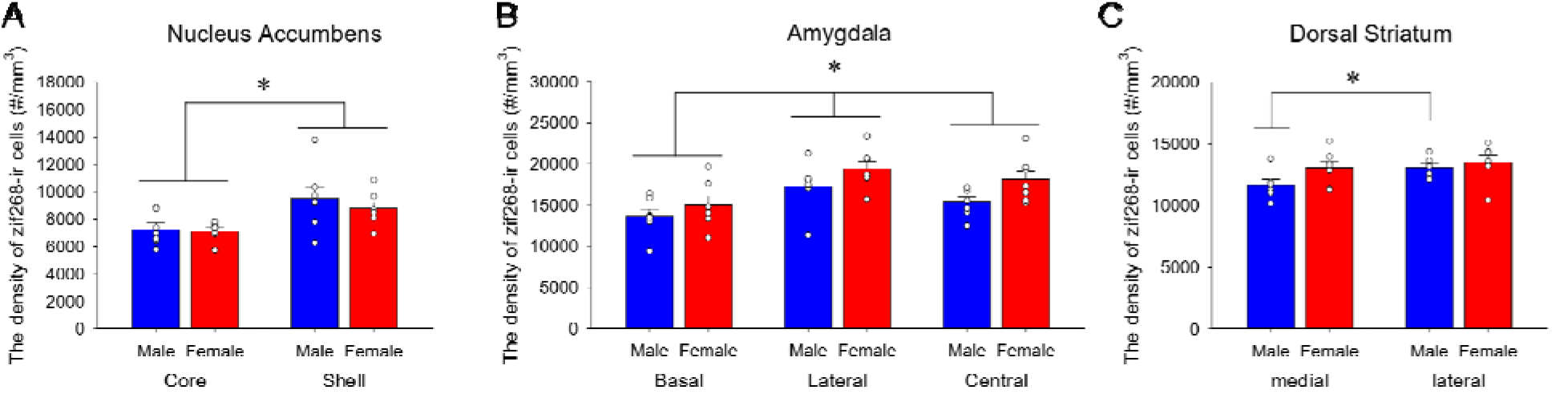
A-C: Mean (±SEM) density of zif268-ir cells in the nucleus accumbens (A), the amygdala (B) and the dorsal striatum (C). The density of zif268-ir cells in the nucleus accumbens shell is greater compared to the nucleus accumbens core (A). Females, compared to males, showed greater density of zif268-ir cells in the amygdala (B). A priori we found that the density of zif268-ir cells was significantly greater in the lDS compared to mDS in males, but not in females. * indicates p < 0.05.

#### Supplemental Results

##### Inter-regional Correlations

In males, within the hippocampus, there were significant correlations between activation of new neurons and activation of subregions of the hippocampus between vDG IdU/zif268-ir cells and vCA3 zif268-ir cells [r(7) = 0.801, p = 0.030], between dDG CldU/zif268-ir cells and dDG zif268-ir cells [r(5) = 0.9926, p = 0.001], and between vDG CldU/zif268-ir cells and dCA3 zif268-ir cells [r(5) = −0.915, p = 0.029]. In females, within the hippocampus, there were significant correlations between the density of dDG IdU/zif268-ir cells and either the vDG DCX/zif268-ir cells [r(7) = 0.848, p = 0.016] or the dCA3 zif268-ir cells [r(7) = −0.794, p = 0.033] and between the density of dDG DCX/zif268-ir cells and the vDG zif268-ir cells [r(8) = −0.745, p = 0.034]. Within females there were also significant correlations between activated new neurons and brain regions outside of the hippocampus (that did not exist in males) between dDG DCX/zif268-ir cells and BLA zif268-ir cells [r(7) = 0.860, p = 0.013] and between vDG IdU/zif268-ir cells and LA zif268-ir cells [r(6) = −0.936, p = 0.006].

Correlations between the 15 different brain regions in males were between the BLA and LA [r(7) = 0.922, p = 0.003], between BLA and mDS [r(7) = 0.789, p = 0.035], between CeA and LA [r(7) = 0.780, p = 0.039] between CeA and IL [r(7) = 0.779, p = 0.039], and between NAc and NAs [r(7) = 0.755, p = 0.050]. In females, between BLA and CeA [r(7) = 0.860, p = 0.013], between BLA and vDG [r(7) = −0.785, p = 0.037], between lDS and NAc [r(7) = 0.910, p = 0.004], between lDS and ACC [r(7) = 0.939, p = 0.002], between lDS and IL [r(7) = −0.760, p = 0.047], between ACC and NAc [r(7) = 0.953, p = 0.001], between PL and dCA1 [r(7) = 0.888, p = 0.008], between dDG and vDG [r(8) = 0.937, p = 0.001], between dDG and dCA1 [r(8) = 0.718, p = 0.045], and between dCA3 and dCA1 [r(8) = 0.790, p = 0.020].

##### Sex differences in inter-regional correlations

The Fischer z-test statistic revealed significant sex differences in correlations between activation of new neurons and activation of subregions of the hippocampus between the density of vDG IdU/zif268-ir cells and vDG DCX/zif268-ir cells [Z(13) = 1.652, p = 0.049], between dDG DCX/zif268-ir cells and vDG zif268-ir cells [Z (13) = 1.723, p = 0.042], vDG DCX/zif268-ir cells and dDG zif268-ir cells [Z(14) = 2.427, p = 0.008], vDG DCX/zif268-ir cells and vDG zif268-ir cells [Z(14) = 2.061, p = 0.020], vDG DCX/zif268-ir cells and vCA1 zif268-ir cells [Z(14) = 1.995, p = 0.023], vDG IdU/zif268-ir cells and vCA3 zif268-ir cells [Z(13) = 1.669, p = 0.048], and dDG CldU/zif268-ir cells and dDG zif268-ir cells [Z(11) = 3.276, p = 0.001]. In addition, there were also significant correlations between activated new neurons and brain regions outside of the hippocampus, between dDG IdU/zif268-ir cells and LA zif268-ir cells [Z(12) = 1.715, p = 0.043], vDG IdU/zif268-ir cells and LA zif268-ir cells [Z(12) = 2.873, p = 0.002].

Furthermore, there were significant sex differences in the correlations between the 15 different brain regions including between LA and BLA [Z(13) = 1.724, p = 0.042], BLA and NAc [Z(13) = 1.765, p = 0.039], lDS and NAc [Z(13) = 2.177, p = 0.015], mDS and NAs [Z(13) = 1.701, p = 0.044], lDS and ACC [Z(13) = 2.696, p = 0.004], NAc and ACC [Z (13) = 2.089, p = 0.018], PL and IL [Z(13) = 2.184, p = 0.014], vDG and BLA [Z(13) = 2.115, p = 0.017], vDG and CeA [Z(13) = 2.133, p = 0.016], vDG and IL [Z(13) = 1.651, p = 0.049], vCA3 and IL [Z(13) = 1.907, p = 0.028], dCA1 and IL [Z (13) = 2.143, p = 0.016], dCA3 and dCA1 [Z (14) = 2.202, p = 0.014], CeA and vCA1 [z(13) = 1.748, p = 0.040], lDS and vCA1 [Z(13) = 1.712, p = 0.043], NAc and vCA1 [Z(13) = 1.967, p = 0.025], IL and vCA1 [Z(13) = 1.881, p = 0.030].

## Notes

### Competing Interest Statement

The authors have declared no competing interest.

